# Bacteriophage density influences the rate of resistance evolution

**DOI:** 10.1101/2025.10.05.680490

**Authors:** Tuan Phan, Anuja Shrestha, Jacob Schow, Craig R. Miller, Tracey L. Peters, James T. Van Leuven

## Abstract

The antagonistic relationship between bacteria and bacteriophages (phages) drives genetic changes that result in phage resistance. Phage resistance mutations arise in natural microbial communities and during the treatment of diseases with phages (phage therapy), making it important to understand the dynamics of resistance acquisition. It is well-established that when bacteria are challenged with phages in dense liquid cultures, bacterial populations quickly become dominated by phage-resistant variants. However, these conditions—well-mixed liquid cultures with high phage concentrations—are not necessarily common in microbial ecosystems. We developed a simplified mathematical model of phage resistance evolution to explore how phage and host concentration impact the dynamics of resistance evolution. The model was parameterized with microbial growth data from two pathogens and their phages: *Pseudomonas aeruginosa* and *Paenibacillus larvae*. Our analyses revealed two fundamental discoveries about resistance evolution. First, phage resistance evolution is predictably governed by a core set of parameters that exhibited high resolution across all bacterial strains: intrinsic growth rate of susceptible bacteria, resistance acquisition rate, fitness cost of resistance, and phage adsorption rate. Second, competitive interactions and fitness costs are the primary drivers of resistance patterns rather than intrinsic mutation rates. We observed three distinct growth patterns—delayed growth, two phase growth, and complete suppression—corresponding to specific parameter regimes as initial phage concentration increased. Two phase growth patterns emerged when competitive dynamics remained balanced, enabling coexistence of susceptible and resistant populations. Complete suppression patterns occurred when low proliferation thresholds combined with extreme competitive asymmetries created unsustainable conditions for resistant bacteria. These findings demonstrate that phage resistance evolution is fundamentally an ecological process where competitive context determines outcomes independently of mutation capacity, with important implications for phage therapy design.

## 1 Introduction

Generalized Lotka-Volterra (gLV) ordinary differential equations (ODEs) are commonly used to describe bacteria-phage dynamics [1, 2, 3]. These ODEs have been modified to accommodate various properties of microbial communities (e.g., carrying capacity, lag in phage growth, phage and bacterial evolution, etc.). The summation of decades of work on this subject is a generally accepted theoretical framework that is mostly consistent with empirical studies [1, 4, 5]. Co-cultured phages and hosts undergo an initial period of dynamics where logistic growth of the host is followed by growth of phage and a crash in host density. The host population can recover and achieve a steady-state (stable coexistence of phage and hosts) when susceptible hosts are at low enough density that they escape phage predation or when phage-resistant variants arise and induce a reciprocal cycle of co-evolution between phage and bacteria. Costs associated with the adaptation of phage resistance may also be invoked to explain why resistant hosts do not entirely displace susceptible hosts [1, 6].

Several intriguing inconsistencies remain between this standard model and empirical results. One pertains to the initial phage inoculum size. Kysela (2007) developed an ODE model to investigate the role of phage mutation rates on the outcome of phage therapy [7]. They found that, contrary to expectation, the peak density of phage-resistant bacteria increases with the multiplicity of infection (MOI) up to about 10-100 phage/host cell, at which point it decreases. The authors suggest that a high MOI decreases the susceptible host density and thus limits the emergence of a second class of phages, which can infect phage-resistant bacteria. This limitation on the phage growth allows the total bacterial density to be higher than if the MOI were lower than around 10. Exploring the impact of phage density on resistance evolution is important because the gut microbiome of most animals is known to be dynamic [8, 9, 10, 11] yet many previous empirical studies explore a fairly narrow range of host and phage densities. Moreover, the pharmacokinetics of phage therapy treatments is still poorly defined, thus it is likely that a wide range of phage and host densities are being unwittingly used. It is imperative to understand the importance of these parameters in the outcome of resistance evolution.

We were initially motivated to model resistance evolution to better understand resistance that was arising while performing Appelmans protocol [12]. Appelmans is a method usually used to expand the host range of a phage cocktail. In the protocol, phages are serially passaged on multiple hosts. Combining phage populations grown on different hosts between passages is supposed to promote phage recombination and rapid adaptation to new hosts. While performing Appelmans in microtiter plates that measures cell density every 10 minutes, we frequently observed cell growth followed by a decrease in cell density, followed by another increase to stationary phase (an expected result given the rise of a phage-resistant population of host cells). However, the standard Appelmans protocol includes a series of 10-fold dilutions of phage, and in these dilution series we often observed that higher concentrations of phage resulted in a more rapid rise in the phage-resistant host population. The expectation from developing and applying mathematical models to these data was that this delay would be explained by the simple mathematics of phage and host replication rates. Instead, we find that simulations using models parameterized off of real growth data do not reflect experimental data. We propose several possible explanations for these disparities based on the relevant literature on phage-bacteria interactions.

## 2 Methods

### 2.1 Bacterial strains and phages

The strains of *Paenibacillus larvae* from ERIC group I were used in the study. Strain Y-3650 was acquired from The University of Nevada Las Vegas [13], whereas the other strains: NRRL B-2605 and ATCC-25747 were received from the American Type Culture Collection (ATCC).

The phage strains specific to *Paenibacillus:* Fern, Xenia, and Scottie were also acquired from the University of Nevada Las Vegas [14]. Phages Fern (KT361649) and Xenia (KT361652) were isolated from infected larvae [15] whereas phage Scottie (MH460825) was isolated from commercial hand cream purchased in the Las Vegas area [16]. These isolates were resequenced when we acquired them. Genome differences were described in Spencer et al. (2025) [17] and the phages were renamed Fern IDv1, Xeni IDv1, and Scot IDv1 to conform with International Committee on the Taxonomy of Viruses (ICTV) standards and differentiate them from the originally published phages which were quite diverged in nucleotide sequence.

All *P. aeruginosa* bacterial strains and their hosts were previously described in [18]. Strains PAK wt Lee, PA103 wt, and PDO300 wt Parsek were received from Dr. Pradeep Sing at the University of Iowa in 2006. PAO1 wt and PA14 wt were a gift from Dr. Jim J. Bull at the University of Idaho, originally given to him from Dr. Joe A. Fralick at Texas Tech University.

*P. aeruginosa* phages included M6, F116, and JM2. M6 was received from Denise Tremblay from the University of Laval Oral Ecology Research Group in 2006. Phages M6 and F116 are commonly used laboratory strains. JM2 was isolated by Jack Milstein at the University of Idaho in 2006 from a Moscow, ID sewer sample. All bacteria and phages were stored at −80 °C in 750 uL of LB media with 30% glycerol. See [18] for a complete description of these phages.

### 2.2 Bacteria and phage propagation

*P. larvae* bacterial strains were streaked from their respective −80°C glycerol stocks onto 1 mg/L of thiamine hydrochloride Brain Heart Infusion (BHI) agar (Catalog#DF0418-17-7, BD Difco™, USA) plates and incubated at 37°C with 5% CO_2_ for 72 hours. A single clean colony from each streaked plate was grown in 1 mg/L of thiamine hydrochloride BHI broth (Catalog#CM1135B, Thermo Scientific™, USA) at 37°C in a shaking incubator at 120 rpm (3 replicates/bacterial strain). After 24 hours of incubation, the bacterial cultures were used for assays. The same bacterial cultures were used to count CFU/mL.

The same basic cell culture methods were used for *P. aeruginosa* except that the propagation media used was Lysogeny broth (LB) and LB-agar. LB is composed of 10 g/L tryptone, 5 g/L yeast extract, and 10 g/L NaCl. Agar was used at a 1.5% with a 0.7% top-agar overlay. *P. aeruginosa* was cultured at 37°C under atmospheric conditions for 14-16 hours.

To resurrect phages from −80 stocks, a loopful of each phage freezer stock was suspended in 200 uL of broth and mixed thoroughly. This stock was serially diluted and titered on *P. larvae* bacterial strain Y-3650, a host permissive to all *P. larvae* phages used in this study. A clean plaque from the Y-3650 titer plate was picked using a sterile cut pipette tip and deposited in a microcentrifuge tube with 750 uL of BHI broth. After gently mixing, 50 uL of chloroform was added to the tube, vortexed thoroughly followed by centrifugation at 15,000 x *g* for 5 minutes. 500 uL of the supernatant was collected and stored in a new tube. This phage stock was then used to make a high titer working phage stock. For this, a 1 mL overnight culture of Y-3650 was prepared from a streaked plate. The 1 mL overnight culture was used to inoculate 9mL of BHI broth. The positive control was a flask of overnight Y-3650 culture: BHI broth (1:10) and the negative control was a flask with 10 mL of BHI broth. Once the OD 600 reached 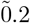(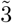 hours), 100 uL of the phage stock was added to the flask. These flasks were incubated in a shaker incubator for 24 hours at 200 rpm. The lysed bacterial cells and spent culture media were aliquoted into 1.5 mL tubes (1 mL/tube). 50 uL of chloroform was added to each tube, vortexed thoroughly and centrifuged at 15,000 x *g* rpm for 5 minutes to fully separate the aqueous layer from the chloroform and any other cell debris. The supernatant was extracted and used as the working phage stock.

Phages were titered by soft-agar overlay. Briefly, a serial dilution of each phage strain was prepared using liquid broth. 3 mL of nutrient-rich top agar (0.7%) enriched with 1mM CaCl_2_ and 1 mM MgCl_2_ was mixed with 100 uL of each host strain and 100 uL of each diluted phage stock, mixed well by briefly vortexing, and poured on the top of a 1.5% agar plate. Plates were then incubated with their lids facing down, at 37°C for 24 hours. The plaques on the plates were counted and the phage stock titers were calculated based on the number of plaques present on the plate and the phage dilutions stock used for plating.

### 2.3 Host range determination

Spot test assays were conducted to determine whether the *P. larvae* specific phages were able to infect the *P. larvae* strains. For this, gridded lines/boxes were made on BHI agar plates and each box was labeled with each phage titration. Then, 100mL of overnight *P. larvae* strain culture was added to a 3 mL of BHI top agar enriched with 1 mM CaCl_2_ (Catalog#CAS 10035-04-8, Fisher Scientific™, USA) and 1 mM MgCl_2_ (Catalog#CAS 7791-18-6, Fisher Scientific, USA) and poured on the labeled BHI agar plate and let it sit down for ~ 30 minutes. The use of CaCl_2_ and MgCl_2_ helps in facilitating phage attachment to their host [13]. Then, 10 uL of serially diluted phage stock (10^0^ to 10^-9^) was spotted onto each labeled gridded box. The plates were left to sit down for about an hour to avoid spreading the phage droplets, followed by incubating plates with lids upright at 37°C with 5% CO_2_ for 24 hours and lysis of each phage activity to its host was noted. If plaques were observed on the plates, then, phage titration plates were made as described below. No plaques on the plates mean that specific *P. larvae* phage failed to lyse that specific *P. larvae* strain on the lawn of which the spots were made. Here, *P. larvae* phage Xenia IDv1 failed to lyse *P. larvae* strain ATCC-25747.

### 2.4 Appelmans protocol

A phage stock was prepared by propagating a single plaque on a permissive host overnight. The resulting lysate was treated with 10% choloroform, then centrifuged at 15,000 x *g* for 5 minutes. 500 mL of the supernatant was collected and stored in a new tube. This phage stock was stored at 4°C during the experiment.

Overnight cultures of *P. larvae* strains were used for the assays. Each assay plate had a total of 8 treatments with 3 replicates per treatment. Of the 8 treatments, 3 were controls: BHI broth control, bacteria control, and phage control. The remaining 5 were the treatments with different phage titers with the same volume and same concentration of bacterial strain. Each assay plate was incubated in a BioTek microplate reader (BioTek^O`^Instrument, Inc. USA) with shaking for 48 hours with the reading of optical density (OD_600_) recorded from each well at the intervals of every 15 minutes.

Overnight cultures of *P. aeruginosa* strains were normalized to an OD 600 nm of 0.3, then diluted 1:10. 1 uL of the diluted hosts was pipetted into each well (a total of roughly 1000 cells per well). Starting with phage stock at a concentration of 1E7 PFU/mL, a dilution series was made from 1E0 to 1E9 dilution and 100 uL of these diluted phages were pipetted into each well. The final volume of the culture was 200 uL per well in 96 well plates. Thus, the maximum resulting MOI was around 1000 PFU/CFU and the minimum MOI was around 1E-6 PFU/CFU.

Cell densities of *P. larvae* and *P. aeruginosa* were measured by counting colonies on agar plates. 2 mL of mBHI was in 24 well plates was inoculated with *P. larvae* Y-3650 at a density of approximately 4.2E6 CFU/mL and 25747 was at a density of 2.4E6 CFU/mL. Phages Fern IDv1, Xeni IDv1, and Scot IDv1 were introduced at a maximum concentration of 2.0E2 PFU/mL, 5.6E2 PFU/mL, and 4.3E2 PFU/mL, respectively.

### 2.5 Mathematical model setup

Adapted from the models used in [4, 5, 19], we propose a simplified mathematical model that describes the interaction among two types of bacterial strains (susceptible and resistant) and one phage strain using generalized Lotka-Volterra equations (Figure 1). We denote by *S*(*t*) and *R*(*t*) (CFU/mL or viable cells/mL), respectively, the population density of the phage-susceptible bacterial strain and the corresponding phage-resistant bacterial strain, and by *P* (*t*) (PFU/mL or viable phage/mL) the population density of the phage strain. In this model, we make the following assumptions. Phage-susceptible strain *S*(*t*) follows the logistic growth with carrying capacity *K* and intrinsic growth rate *r*. This strain can acquire phage resistance through genetic materials transferred from the phage-resistant strain *R*(*t*) at the rate of δ. After transitioning from susceptible to resistant type, the phage-resistant strain is subject to a fitness cost *γ*, lowering its intrinsic growth rate. When attaching to susceptible strain *S*(*t*), the phage strain can adsorb the susceptible at the rate of *ϕ*. After the lysis of the susceptible, the phage will burst out with the rate of *β*. The phage decays at the rate of *m*. The dynamics of the densities of two types of bacterial strains and the phage strain are depicted by the following system of 3 ordinary differential equations.

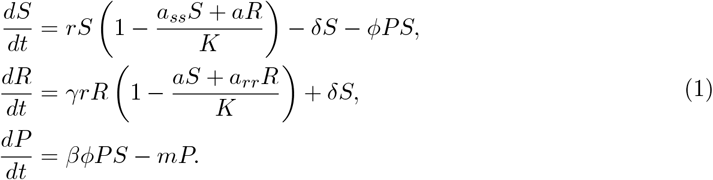

**Figure 1.**
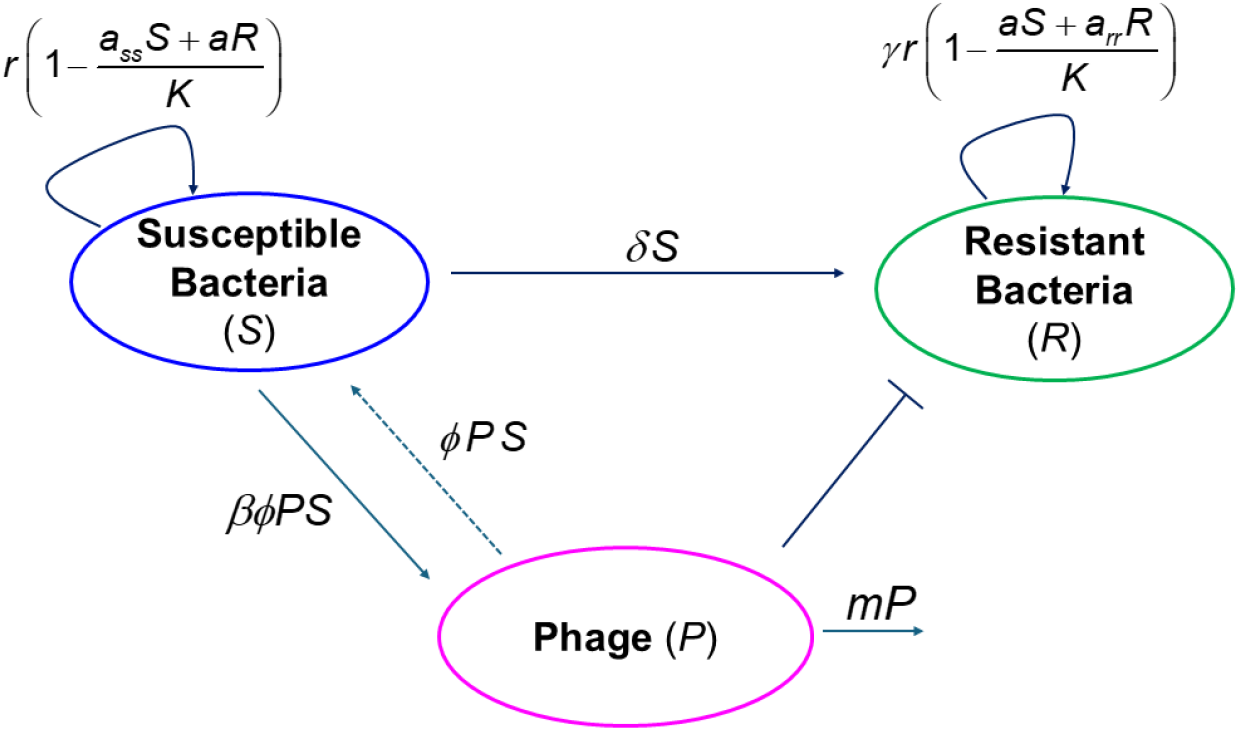
Schematic of the model. Parameters of the model are depicted in Table 1.

## 3 Results

### 3.1 MOI dramatically impacts the dynamics of resistance evolution

To measure how the multiplicity of infection (MOI) impacts resistance evolution, a series of growth experiments were performed on a shaken, incubating microplate reader. We chose eight different hosts (5 strains of *P. aeruginosa* and 3 strains of *P. larvae*) with four different phage challenges (Table 2). These combinations were chosen because these challenges always resulted in phage-resistant colonies overnight when tested on solid agar plates [17, 18]. Unexpectedly, for most (8/14) challenges, we did not observe resistance evolution in liquid culture at any dilution of phage (Table 2). We considered resistance evolution to have occurred when bacterial growth was delayed compared to control or when two periods of bacterial growth were measured (Figure 2). These two phases of growth included an immediate initial growth period that matched the no-phage control that was followed by bacterial death (drop in cell density) and then a rebound in bacterial growth (presumably the rise of a resistant variant(s), Figure 2). The phage-host combinations where we observed these dynamics are labeled “Two phage growth” in (Table 2). The *P. aeruginosa* experiments were initially designed to expand the host range of a phage cocktail through experimental evolution (Appelmans protocol). Thus, *P. aeruginosa* strains were challenged with a mix of three phages. We observed two-phase growth dynamics in strains PA103 and PAK. Strains PA14 and PA01 also grew after a delay, indicating resistant strains evolved. For *P. larvae*, we did not observe any strains with only delayed growth, only two-phase growth. Therefore, we chose to use only *P. aeruginosa* strains with this two-phase growth in our modeling. The *P. larvae* experiments were performed with one phage strain at a time.

**Table 1:** Parameters and their ranges for the system (1)

| Parameters | Description | Values | Units | References |
| --- | --- | --- | --- | --- |
| $r$ | intrinsic growth rate of $S$ | [0.462, 1.65] | $\text{h}^{-1}$ | [20] |
| $a$ | interspecific competition rate between $S$ and $R$ | | dimensionless | |
| $a_{ss}$ | intraspecific competition rate of $S$ with itself | | dimensionless | |
| $a_{rr}$ | intraspecific competition rate of $R$ with itself | | dimensionless | |
| $K$ | system-wide carrying capacity | $10^9$ | CFU/mL | [21] |
| $\gamma$ | fitness cost of phage resistance | [0, 1] | dimensionless | [4] |
| $\delta$ | rate of resistance acquisition | $[10^{-6}, 10^{-3}]$ | $\text{h}^{-1}$ | [4] |
| $\phi$ | adsorption rate of phage $P$ when attaching to bacteria $S$ | $[10^{-8}, 10^{-6}]$ | $\text{mL} \cdot \text{PFU}^{-1} \cdot \text{h}^{-1}$ | [1, 4, 19] |
| $\beta$ | the burst size of phage $P$ | [10, 100] | $\text{PFU} \cdot \text{CFU}^{-1}$ | [1, 4] |
| $m$ | decay rate of phage $P$ | $[1.5 \cdot 10^{-3}, 0.1]$ | $\text{h}^{-1}$ | [22] |

**Table 2:** Results of growth experiments using 8 hosts with 4 different phages.

| Strains | Fern_IDv1 | Scot_IDv1 | Xeni_IDv1 | Cocktail |
| --- | --- | --- | --- | --- |
| *P. larvae* 3650 | **Two phase growth** | Complete suppression | Complete suppression | Not tested |
| *P. larvae* 25747 | **Two phase growth** | Complete suppression | Complete suppression | Not tested |
| *P. larvae* 2605 | Complete suppression | Complete suppression | Complete suppression | Not tested |
| *P. aeruginosa* PA103 | Not tested | Not tested | Not tested | **Two phase growth** |
| *P. aeruginosa* PA14 | Not tested | Not tested | Not tested | Delayed growth |
| *P. aeruginosa* PA01 | Not tested | Not tested | Not tested | Delayed growth |
| *P. aeruginosa* PAK | Not tested | Not tested | Not tested | **Two phase growth** |
| *P. aeruginosa* PDO300 | Not tested | Not tested | Not tested | Complete suppression |

**Figure 2.**
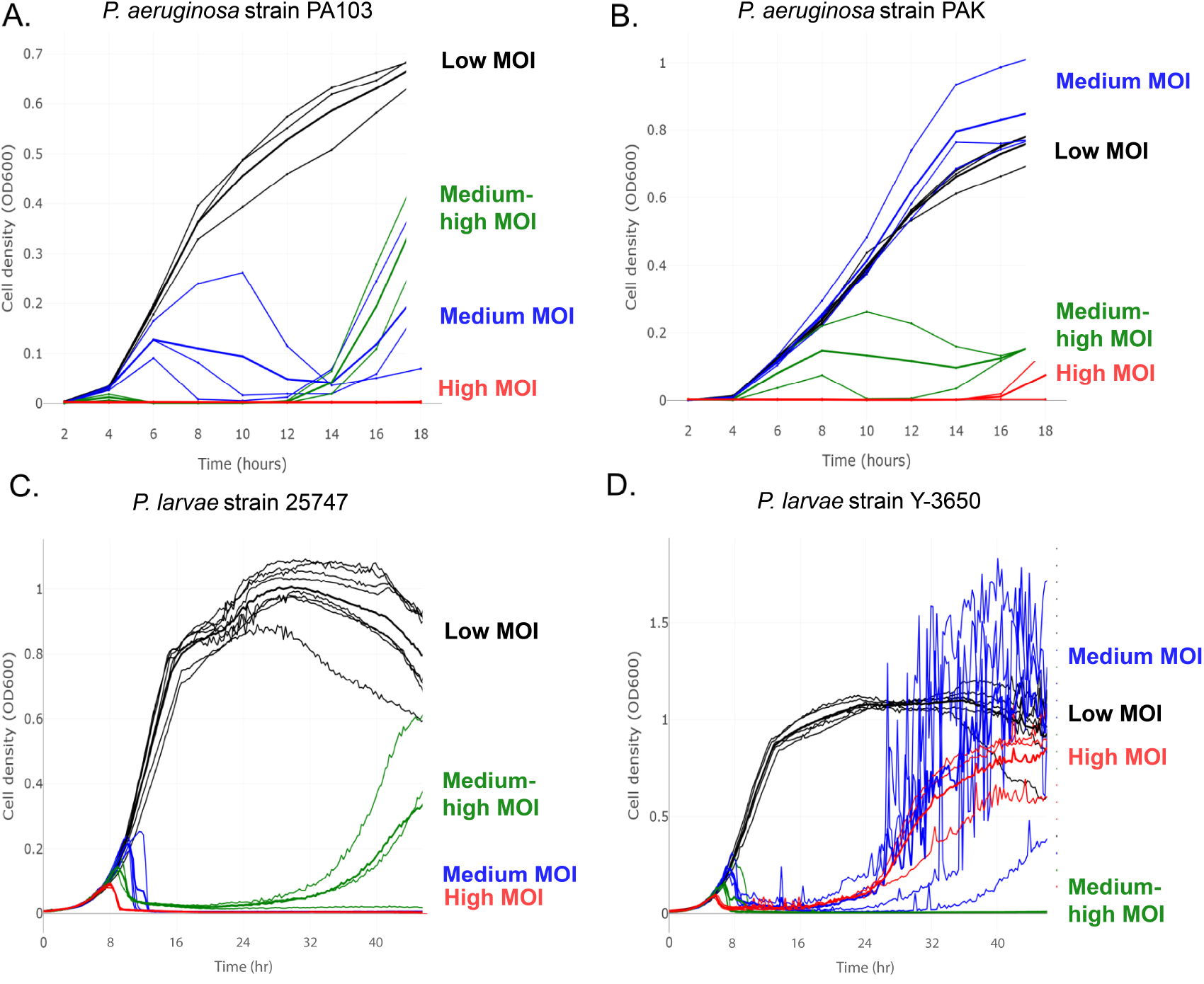
Dynamics of growth experimental data of four host strains that exhibit two-phase growth patterns. Panels A and B, respectively, show growth dynamics of *P. aeruginosa* PA103 and PAK at round 3 of Appelmans protocol with different dilutions of a cocktail of phages. Panels C and D, respectively, represent growth dynamics of *P. larvae* 25747 and *P. larvae* Y-3650 with different dilutions of phage Fern IDv1 (3 replicates for each dilution).

### 3.2 Model parameterization

In this research, we will use the mathematical model (1) to test our central hypothesis that initial phage density influences the dynamics of resistance evolution. To do so, we aim to fit the model (1) to experimental growth data of four host strains *P. aeruginosa* PA103, *P. aeruginosa* PAK, *P. larvae* 3650 and *P. larvae* 25747 as depicted in Table 2 and Figure 2. We selected these host strains because phage-resistant colonies were always observed on solid agar plates. To examine resistance evolution in datasets of these four host strains using our mathematical model, we classified their experimental growth data into four groups (Low, Medium, Medium High, High) based on the initial MOI, as shown in Figure 2. For example, in Panel A, the growth data for PA103 strain under three conditions—no phage control (Ctrl) and two very low concentration phage treatments (10^−9^ and 10^−8^ dilutions of the starting phage stock)–were grouped into a low phage treatment group (Low) because all three exhibited similar growth patterns with minimal to no resistance. Here, the Low data were calculated by averaging the data from the no-phage, 10^−9^, and 10^−8^ dilutions. Next, we grouped three phage treatments (10^−7^, 10^−6^, and 10^−5^ dilutions) into the Medium group since they exhibited a two-phase growth pattern, two phage treatments (10^−4^, 10^−3^ dilutions) into the Medium High group as they showed delayed growth pattern, and three phage treatments (10^−2^ dilution, 10^−1^ dilution, and undiluted phage stock) into the High group since complete suppression pattern was observed.

Next, we examined the dynamics of the mathematical model (1) to identify key parameters related to the dynamics of resistance evolution. By the first equation of (1), phage susceptible bacterial population *S* does not grow or decay when 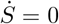, which is equivalent to 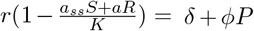. The term 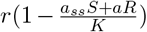 represents the effective growth rate of phage susceptible bacteria, while *ϕP* represents the likelihood that a phage susceptible bacterium is absorbed by phages and δ stands for the likelihood that a phage susceptible bacterium mutates into phage resistant type. When phage concentration *P* is abundant such that 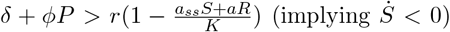, phages can absorb susceptible bacteria quickly, causing the population of susceptible bacteria to decline. Conversely, when phage concentration *P* is low such that 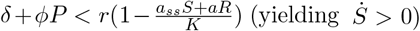, the effective growth rate of susceptible bacteria dominates the likelihood of mutation and adsorption by phage, allowing the susceptible bacterial population to grow.

According to the second equation of (1), phage resistant bacterial population *R* grows (or decays) when 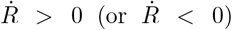, which is equivalent to 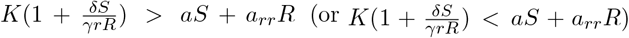. The parameter *a*_*rr*_ represents the intraspecific competition rate of the phage resistant bacteria *R* with itself, quantifying how strongly resistant bacteria compete against other resistant bacteria within the same population for limited resources. Similarly, the parameter *a* represents the interspecific competition rate between susceptible (*S*) and resistant (*R*) bacteria, indicating how strongly the susceptible bacteria inhibit the growth of resistant bacteria. When one of these two parameters or both are high such that 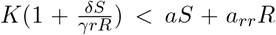, the resistant bacterial population *R* decays. Biologically, this indicates that resistant bacteria’s growth is significantly limited by competition—both from other resistant bacteria and from the susceptible population—even at relatively low density, as they rapidly compete for available resources or occupy niches, quickly saturating the environment and preventing further population growth. Conversely, when both *a* and *a*_*rr*_ are low (indicating weak competition both within resistant strain and between strains) such that 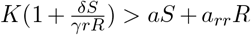, the population *R* grows because resistant bacteria face less competition, allowing them to coexist more easily and reach higher density. The relative magnitudes of these two competition parameters also influence the system’s dynamics: when *a > a*_*rr*_, the presence of susceptible bacteria more strongly inhibits resistant bacteria growth than competition among resistant bacteria themselves, whereas when *a < a*_*rr*_, intraspecific competition becomes the dominant limiting factor. Thus, the values of both *a* and *a*_*rr*_ crucially influence how effectively resistance evolves and spreads, shaping practical strategies in managing bacterial populations through phage-based intervention.

Regarding phage concentration dynamics, the third equation of (1) implies that phage concentration *P* remains constant when 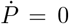, which is equivalent to 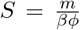. We define this threshold as 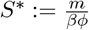, which mathematically represents the critical density of susceptible bacteria required for the phage population to increase. Biologically, this threshold signifies the minimum concentration of susceptible hosts necessary to sustain phage replication and proliferation. When the density of susceptible bacteria exceeds *S*^∗^ (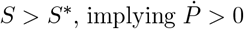), phages can effectively infect hosts, leading to increased phage concentration as each phage infects enough susceptible bacteria to produce sufficient progeny to replace those that decay naturally. Conversely, if *S* falls below *S*^∗^ (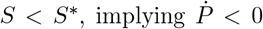), phage replication becomes insufficient to offset phage decay, resulting in declining phage numbers. This concept of a critical host density for phage proliferation is well-documented in phage ecology [23, 24, 25]. We refer to this threshold as the *proliferation threshold* —the minimum susceptible bacterial density required to support effective phage replication.

From fitted results in Table 3, we consistently observed three typical growth patterns—delayed growth, two phase growth, and complete suppression—occurring across all four strains as initial phage concentration increases from Low to High. For strains exhibiting complete suppression patterns (except 25747 at Medium and Y-3650 at Medium High), all required high initial phage concentrations to be suppressed. In these strains, the fitted parameters *a* and *a*_*ss*_ exhibited very high values while the fitted proliferation threshold *S*^∗^ and the fitted value of *γ* showed low values, explaining why both susceptible and resistant populations declined under high phage pressure (Figure 4D, Figure 5D, and Figure 6D). Interestingly, strain 25747 could be completely suppressed even at Medium phage treatment, with the fitted parameter *a*_*rr*_ exhibiting a very high value while the fitted parameter *r* showed a very low value. This indicates that resistant bacteria were totally suppressed, while susceptible bacteria initially grew but then rapidly declined (Figure 6B)—dynamics strikingly similar to those in strain Y-3650 at Medium High (Figure 7C).

**Table 3:** Results of fitted parameters for experimental growth datasets in four strains PA103 Rd3 Cocktail, PAK Rd3 Cocktail, 25747 Fern IDv1 and Y-3650 Fern IDv1 using the mathematical model (1).

| # | Parameter | PA103 Rd3 Cocktail |  |  |  | PAK Rd3 Cocktail |  |  |  |
| --- | --- | --- | --- | --- | --- | --- | --- | --- | --- |
|  |  | Low | Medium | Medium High | High | Low | Medium | Medium High | High |
| 1 | $r$ | 1.0146 | 1.6458 | 1.4538 | 1.0538 | 1.4146 | 1.5146 | 1.6497 | 0.8538 |
| 2 | $a$ | 5.3525 | 2.6901 | 0.1 | 7648.2 | 7.9525 | 8.5525 | 42.19945 | 368989 |
| 3 | $a_{ss}$ | 40.325 | 102.675 | 188.6 | 3425.5 | 38.525 | 39.325 | 83.69450 | 1990774 |
| 4 | $a_{rr}$ | 12.7417 | 0.0011 | 14.3818 | 1.5818 | 12.9417 | 11.5417 | 0.001375 | 0.0818 |
| 5 | $\delta$ | 0.00041 | 0.00041 | 0.0002 | 0.0002 | 0.00041 | 0.00041 | 0.001 | 0.0002 |
| 6 | $\gamma$ | 0.4009 | 0.115 | 0.5274 | 0.2274 | 0.7009 | 0.7009 | 0.50468 | 0.3274 |
| 7 | $\phi$ | $10^{-8}$ | $10^{-8}$ | $10^{-8}$ | $10^{-8}$ | $10^{-8}$ | $10^{-8}$ | $10^{-8}$ | $10^{-8}$ |
| 8 | $\beta$ | 11.525 | 37.8719 | 96.8 | 100 | 11.625 | 10.425 | 10 | 70.7 |
| 9 | $m$ | 0.079 | 0.0478 | 0.0401 | 0.0401 | 0.079 | 0.079 | 0.1 | 0.0401 |
| 10 | $S_0$ | 436680 | 124828 | 0.1 | 185104 | 70477 | 43869 | 34776.56 | 222981 |
| 11 | $R_0$ | 1534810 | 942122 | 383.4 | 84259 | 11995.6 | 9793.4 | 0.03125 | 76573.4 |
| 12 | $P_0$ | 0.8 | 2.0594 | 10 | 999.2 | 1 | 5.2 | 10 | 439 |
| | $S^* = \frac{m}{\beta\phi}$ | 685466 | 126215 | 41426 | 40100 | 679570 | 757794 | 1000000 | 56719 |
|  |  | Delayed growth | 2-phase growth | Delayed growth | Complete suppression | Delayed growth | Delayed growth | 2-phase growth | Complete suppression |

| # | Parameter | 25747 Fern_IDv1 |  |  |  | Y-3650 Fern_IDv1 |  |  |  |
| --- | --- | --- | --- | --- | --- | --- | --- | --- | --- |
|  |  | Low | Medium | Medium High | High | Low | Medium | Medium High | High |
| 1 | $r$ | 0.5146 | 0.5146 | 1.1499 | 1.3499 | 0.7457 | 1.5531 | 0.7790 | 1.1478 |
| 2 | $a$ | 7.4525 | 16.4525 | 0.17 | 780.97 | 4.7349 | 1.52E-7 | 0.2603 | 468.54 |
| 3 | $a_{ss}$ | 12.625 | 9.425 | 89.6945 | 107.2945 | 49.0987 | 42.0266 | 56.4159 | 89.7757 |
| 4 | $a_{rr}$ | 11.7417 | 54078.94 | 8.99 | 136.99 | 9.5935 | 9.0342 | 1023.31 | 12.6446 |
| 5 | $\delta$ | 0.00041 | 0.00041 | 0.00098 | 0.00098 | 0.001 | 0.001 | 0.0008 | 0.001 |
| 6 | $\gamma$ | 0.7009 | 0.0009 | 0.1274 | 0.0274 | 0.8575 | 0.1751 | 0.9957 | 0.3304 |
| 7 | $\phi$ | 1E-9 | 1E-8 | 1E-8 | 1E-8 | 1.1E-8 | 2E-8 | 2E-8 | 5.94E-8 |
| 8 | $\beta$ | 11.375 | 22.575 | 28.4 | 36 | 22.436 | 10.676 | 13.429 | 10.943 |
| 9 | $m$ | 0.079 | 0.079 | 0.037 | 0.037 | 0.1 | 0.096 | 0.0027 | 0.053 |
| 10 | $S_0$ | 124753 | 140219 | 8539 | 36417 | 109472 | 2303 | 403831 | 66084 |
| 11 | $R_0$ | 88492 | 1124436 | 30502 | 248870 | 100032 | 1184 | 134 | 1027 |
| 12 | $P_0$ | 3 | 7.5 | 21 | 2024 | 0.3 | 3.48 | 72.96 | 356.45 |
| | $S^* = \frac{m}{\beta\phi}$ | 6945055 | 349945 | 130282 | 102778 | 441289 | 447589 | 10090 | 81858 |
|  |  | Delayed growth | Complete suppression | 2-phase growth | Complete suppression | Delayed growth | 2-phase growth | Complete suppression | 2-phase growth |

In contrast, for strains exhibiting two-phase growth patterns (PA103 Rd3 at Medium, PAK Rd3 at Medium High, 25747 at Medium High, and Y–3650 at Medium and High), we observed that the intraspecific competition rate *a*_*rr*_ for resistant bacteria was much lower than the intraspecific competition rate *a*_*ss*_ for susceptible bacteria. This suggests that the resistant population *R* initially grew more slowly than the susceptible population *S* but eventually dominated (Figure 4B, Figure 5C, Figure 6C, Figure 7B and Figure 7D). For strains exhibiting delayed growth patterns, the gap |*a*_*ss*_ − *a*_*rr*_| was not too large across all strains, and the fitted proliferation threshold *S*^∗^ was quite high, indicating that phages required considerable time to absorb susceptible bacteria and grow. The resistant population *R* was initially suppressed for 5 to 10 hours before growing dramatically fast, preceding the decline of the susceptible population *S*.

Our observations yielded several interesting results. Complete suppression patterns occasionally occurred at Medium phage treatment (25747 at Medium), which is relatively rare. Two phase growth patterns frequently occurred at either Medium or Medium High phage concentrations across strains, but interestingly, this pattern was also possible at High phage concentrations (Y-3650 at High). These findings suggest that resistance evolution may occur across various phage concentrations—medium, medium high, and even high—making resistance evolution relatively unpredictable.

### 3.3 Sensitivity analysis

Resolution matrices provide a quantitative measure of parameter identifiability in mathematical models fitted to experimental data, with diagonal elements indicating how well each parameter can be uniquely determined and off-diagonal elements revealing correlations between parameters that may confound parameter estimation. In the context of our mathematical model (1), these matrices are essential tools for understanding which biological processes most strongly influence bacterial resistance evolution and which parameters can be reliably estimated from experimental growth curves. The brightness of diagonal elements (approach 1.0) indicates high parameter resolution, while darker values suggest parameters that are poorly constrained by the available data. Results of model resolution matrices generated by fitted parameters across four bacterial strains in Table 3 are shown in Figures 8, 9, 10, and 11. Table 4 summarizes well-resolved parameters of the model (1) based three types of growth pattern (delayed growth, two phase growth, and complete suppression). Our analysis across four bacterial strains under varying phage concentrations revealed distinct patterns of parameter identifiability that correlate with the observed growth dynamics.

**Table 4:** Growth pattern based parameter resolution. Note that the description of parameter indices can be referred in Table 3 and Table 1.

| Pattern | Strains & Panels | Well-resolved parameters (by index) |
| --- | --- | --- |
| Delayed growth | PA103 Rd3 (Fig. 8A, 8C) | 1, 5, 6, 7, 9 |
|  | PAK Rd3 (Fig. 9A, 9B) | 1, 4, 5, 6, 7, 9 |
|  | 25747 (Fig. 10A) | 1, 2, 3, 4, 5, 6, 7, 8, 9, 12 |
|  | Y-3650 (Fig. 11A) | 1, 2, 3, 4, 5, 6, 7, 8, 9, 12 |
| Two phase growth | PA103 Rd3 (Fig. 8B) | 1, 4, 5, 6, 7, 9 |
|  | PAK Rd3 (Fig. 9C) | 1, 5, 6, 7, 9 |
|  | 25747 (Fig. 10C) | 1, 2, 3, 4, 5, 6, 7, 8, 9, 12 |
|  | Y3650 (Fig. 11B, Fig. 11D) | 1, 2, 3, 4, 5, 6, 7, 8, 9, 12 |
| Complete suppression | PA103 Rd3 (Fig. 8D) | 1, 5, 6, 7 |
|  | PAK Rd3 (Fig. 9D) | 1, 5, 6, 7, 10, 11 |
|  | 25747 (Fig. 10B, 10D) | 1, 5, 6, 7, 8, 9, 12 |
|  | Y-3650 (Fig. 11C) | 1, 2, 3, 4, 5, 6, 7, 8, 9, 11, 12 |

In delayed growth patterns, predominantly observed at low phage concentrations, our resolution matrices consistently showed high identifiability for a core set of parameters across all strains. All four strains exhibiting delayed growth (PA103 at Low and Medium High, PAK at Low and Medium, 25747 at Low, and Y-3650 at Low) resolved five fundamental parameters: intrinsic growth rate of susceptible bacteria (*r*), rate of resistance acquisition (δ), fitness cost of resistance (*γ*), phage adsorption rate (*ϕ*), and phage decay rate (*m*) - corresponding to parameters 1, 5, 6, 7, and 9 in Table 4. Additionally, *P. larvae* strains (25747 and Y-3650) showed broader parameter resolution, including all competition coefficients (*a, a*_*ss*_, *a*_*rr*_ - parameters 2, 3, 4), phage burst size (*β* - parameter 8), and initial phage concentration (*P*_0_ - parameter 12). These parameters exhibited strong diagonal dominance with minimal off-diagonal correlations in the resolution matrices (Figures 8A, 8C, 9A, 9B, 10A, 11A), indicating robust parameter identifiability. This pattern suggests that under low phage pressure, bacterial survival depends primarily on intrinsic growth and resistance acquisition mechanisms, while *P. larvae* strains maintain additional sensitivity to competitive dynamics and phage kinetics even at low phage concentrations.

The transition to two-phase growth, characterized by an initial bacterial expansion followed by phage-induced suppression and subsequent resistant cell emergence, revealed varied parameter resolution patterns across strains. PA103 at Medium (Figure 8B) resolved the four core parameters (*r*, δ, *γ, ϕ*) plus phage decay rate (*m*) and notably, the intraspecific competition rate of resistant bacteria (*a*_*rr*_), suggesting that competition within the resistant population becomes important during the transition between growth phases. PAK at Medium High (Figure 9C) showed more limited resolution, identifying only the core parameters and phage decay rate without resolving competition coefficients. In contrast, 25747 at Medium High and Y-3650 at both Medium and High (Figures 10C, 11B, 11D) demonstrated comprehensive parameter identifiability across nearly all model components, including all competition parameters (*a, a*_*ss*_, *a*_*rr*_), phage burst size (*β*), and initial conditions. The resolution of phage burst size exclusively in the *P. larvae* systems reflects the critical role of phage reproductive capacity in determining the depth and duration of the suppression phase between bacterial growth waves. This strain-specific variation in parameter resolution indicates that different bacterial species utilize distinct mechanisms to navigate the complex dynamics of two-phase growth under moderate phage pressure.

Under complete suppression conditions, where high phage concentrations prevented bacterial recovery, our resolution matrices revealed the most complex parameter interaction patterns. The four core parameters (*r*, δ, *γ, ϕ*) remained well-resolved across all strains, but additional parameters achieved high identifiability depending on the bacterial system. PA103 showed the most constrained parameter resolution under suppression (Figure 8D), resolving only the four core parameters. PAK Rd3 exhibited broader resolution at high phage concentration (Figure 9D), additionally resolving initial susceptible and resistant populations (*S*_0_, *R*_0_). In contrast, both *P. larvae* strains maintained comprehensive parameter resolution even under suppression conditions. 25747 resolved nearly all parameters including competition coefficients, phage burst size (*β*), and initial phage concentration (*P*_0_) at both Medium and High suppression conditions (Figures 10B, 10D). Similarly, Y3650 at Medium High showed broad parameter resolution including competition parameters, initial populations, and phage kinetics (Figure 11C). These patterns suggest that different bacterial species employ distinct adaptive strategies under extreme phage pressure, with *P. larvae* strains maintaining more complex regulatory networks that remain identifiable even when populations are suppressed.

Our sensitivity analysis reveals that resistance evolution is fundamentally driven by a core parameter set (*r*, δ, *γ, ϕ*) that shows consistent high resolution across all growth patterns and bacterial strains. These four parameters—intrinsic growth rate of susceptible bacteria, resistance acquisition rate, fitness cost of resistance, and phage adsorption rate—represent the fundamental biological processes governing resistance evolution regardless of phage concentration or bacterial species. Beyond this universal core, parameter resolution patterns diverge based on both growth dynamics and bacterial strain. Phage decay rate (*m*) emerged as consistently important only in delayed growth conditions across all strains, suggesting its critical role when bacteria can outlast phage presence. Competition parameters showed variable resolution: *P. larvae* strains maintained identifiability of competition coefficients across most conditions, while *P. aeruginosa* strains resolved these parameters sporadically—PA103 identifying *a*_*rr*_ during two-phase growth and PAK identifying initial populations under suppression. Notably, phage burst size (*β*) was resolved exclusively in *P. larvae* systems across all growth patterns, indicating fundamental differences in how these species respond to phage reproductive dynamics. The distinct resolution patterns between bacterial species—with *P. larvae* maintaining comprehensive parameter identifiability even under extreme suppression while *P. aeruginosa* shows minimal resolution—demonstrate that therapeutic approaches must be tailored to specific bacterial systems rather than following universal protocols. These findings suggest that while core resistance mechanisms operate universally, the ecological and competitive contexts that ultimately determine treatment outcomes are highly species-specific.

## 4 Discussion

The parameterization of our mathematical model (1) to experimental growth data from four bacterial strains revealed two fundamental discoveries about phage resistance evolution. First, resistance evolution dynamics are predictably governed by a core set of identifiable parameters that consistently exhibited high resolution across all growth patterns and bacterial strains: intrinsic growth rate of susceptible bacteria (*r*), resistance acquisition rate (δ), fitness cost of resistance (*γ*), and phage adsorption rate (*ϕ*). Second, and more importantly, competitive interactions and fitness costs are the primary drivers of resistance patterns rather than intrinsic mutation rates. This result fundamentally reframes resistance evolution from a primarily genetic/mutational process to an ecological/competitive process, with major implications for therapeutic strategies [26].

The consistent identifiability of the core parameter set across diverse bacterial species and phage concentrations (Table 4, Figures 8-11) demonstrates that resistance evolution follows predictable mathematical rules rather than stochastic processes. These parameters exhibited strong diagonal dominance with minimal correlations in resolution matrices, indicating their fundamental biological importance regardless of experimental conditions. Notably, phage-specific factors—including burst size (*β*), decay rate (*m*), and initial phage population (*P*_0_)—gained prominence only under moderate to high phage pressures, suggesting a hierarchical model where basic resistance mechanism operate universally while phage dynamics become increasingly influential as selection pressure intensifies [27, 28].

The critical insight from our parameter dynamics analysis (Figure 3) is that competitive interactions, not mutation rates, determine resistance evolution outcomes. While resistance acquisition rate δ remains relatively stable across phage concentrations in most strains, competition parameters (*a, a*_*rr*_) varied by orders of magnitude, driving completely different growth patterns. This observation demonstrates that the same bacterial strain with nearly identical intrinsic resistance capacity can exhibit fundamentally different evolutionary outcomes—from successful resistance emergence to complete population collapse—depending solely on the competitive landscape established by phage concentration.

**Figure 3.**
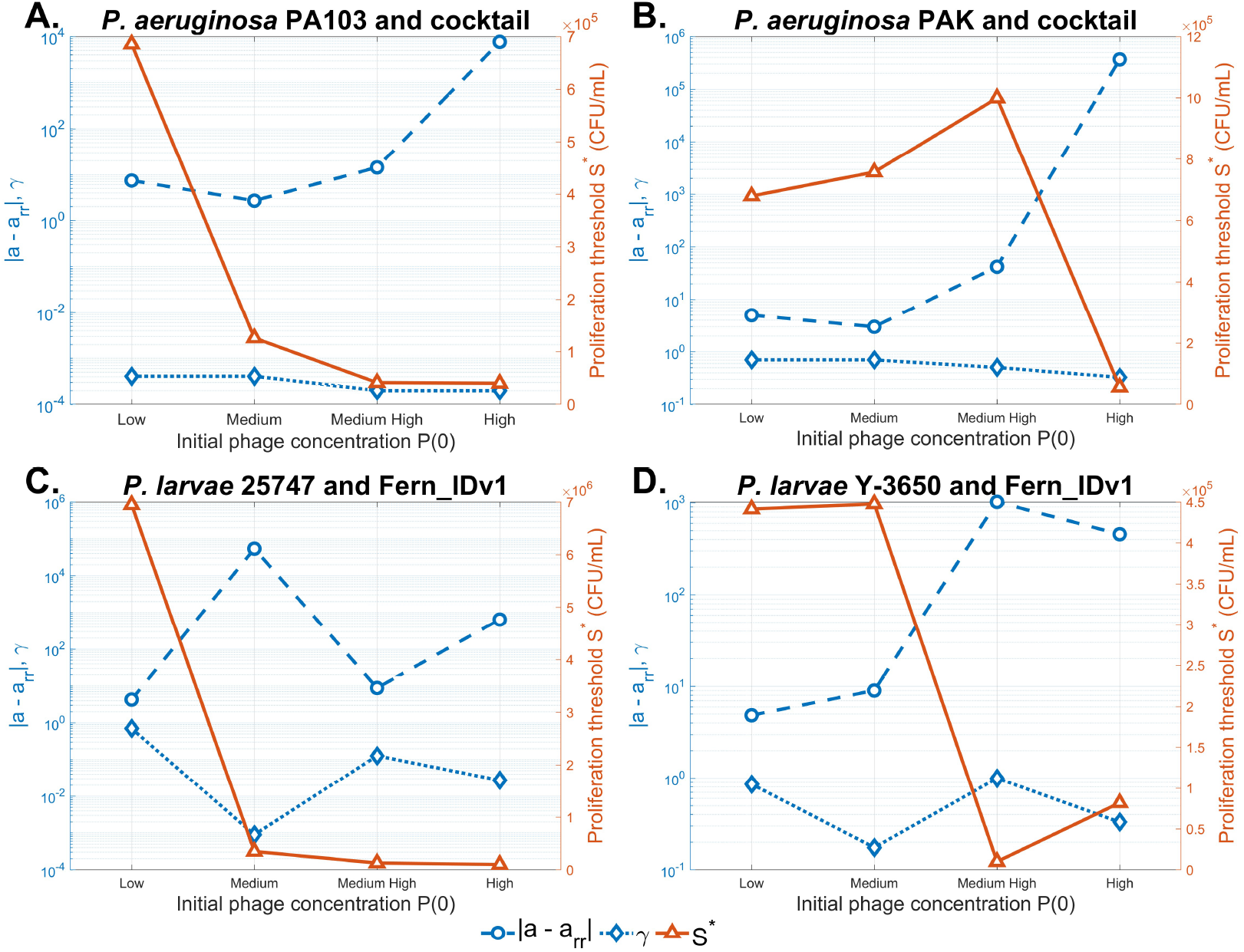
Dynamics of the fitted gap |*a* − *a*_*rr*_| and the fitted parameters *γ, S*^∗^ vs. initial phage concentration *P*_0_ in four host strains treated with a phage cocktail or a single phage.

**Figure 4.**
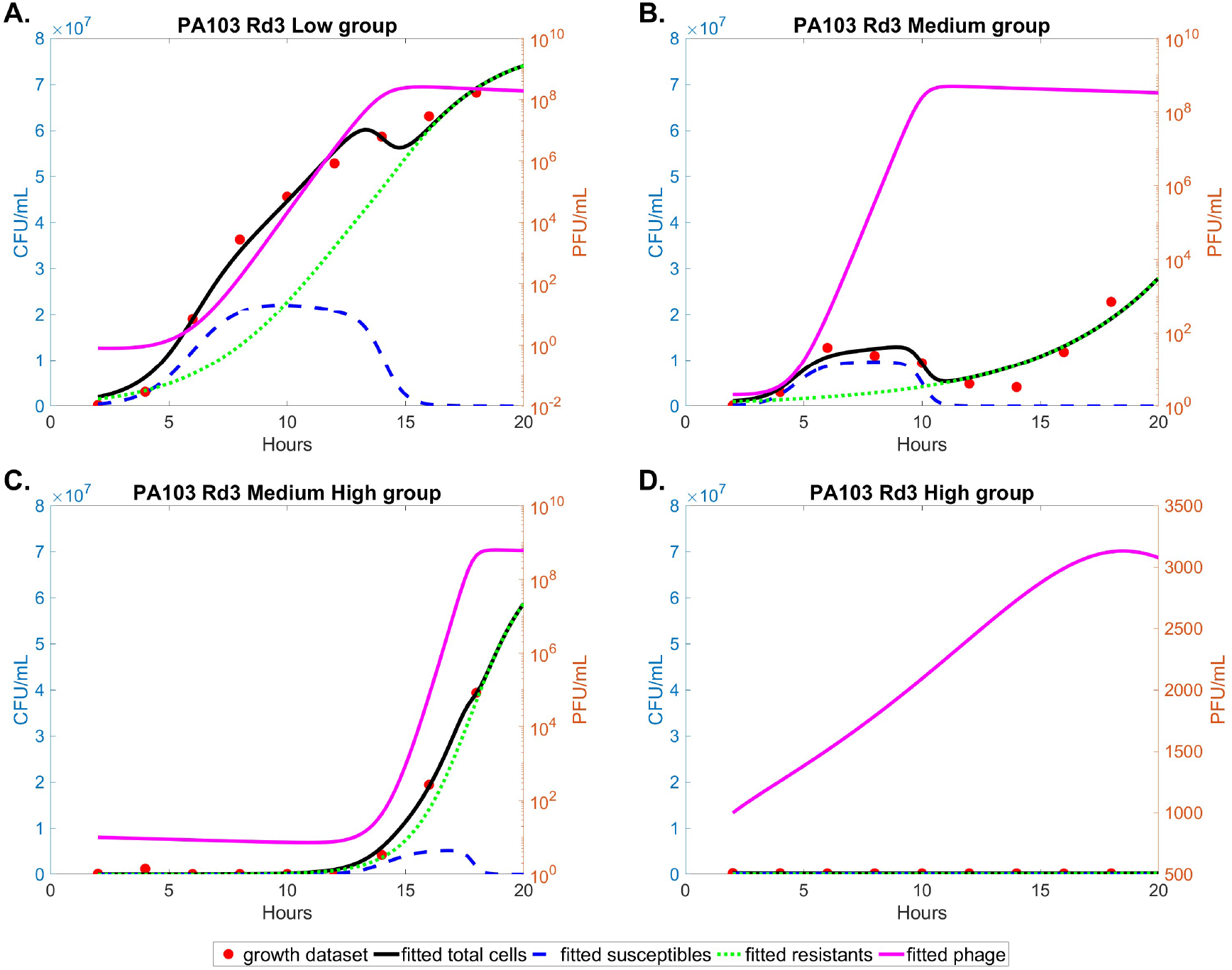
Dynamics of the mathematical model (1) when it is fitted to experimental growth dataset of strain PA103 Rd3.

**Figure 5.**
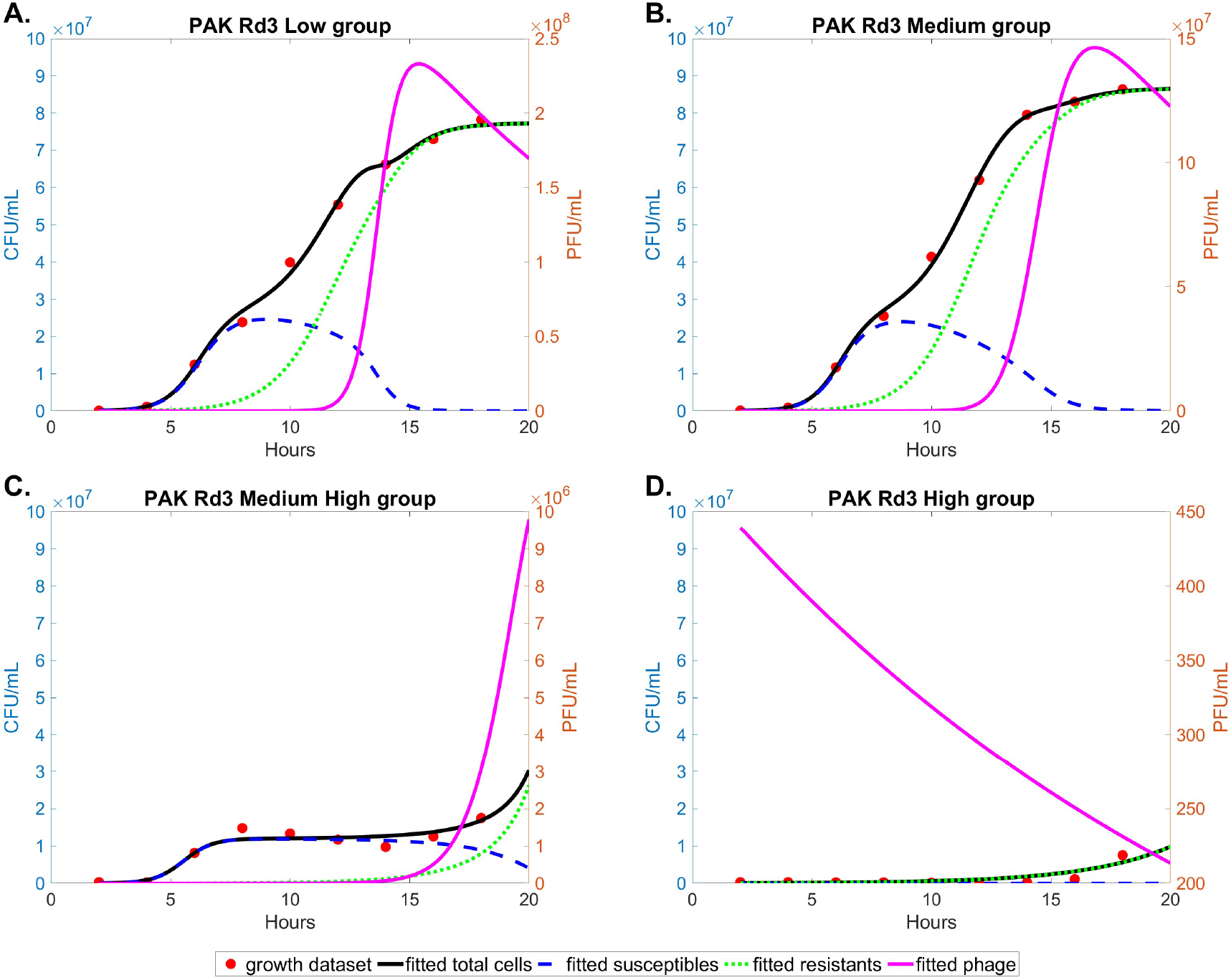
Dynamics of the mathematical model (1) when it is fitted to experimental growth dataset of strain PAK Rd3.

**Figure 6.**
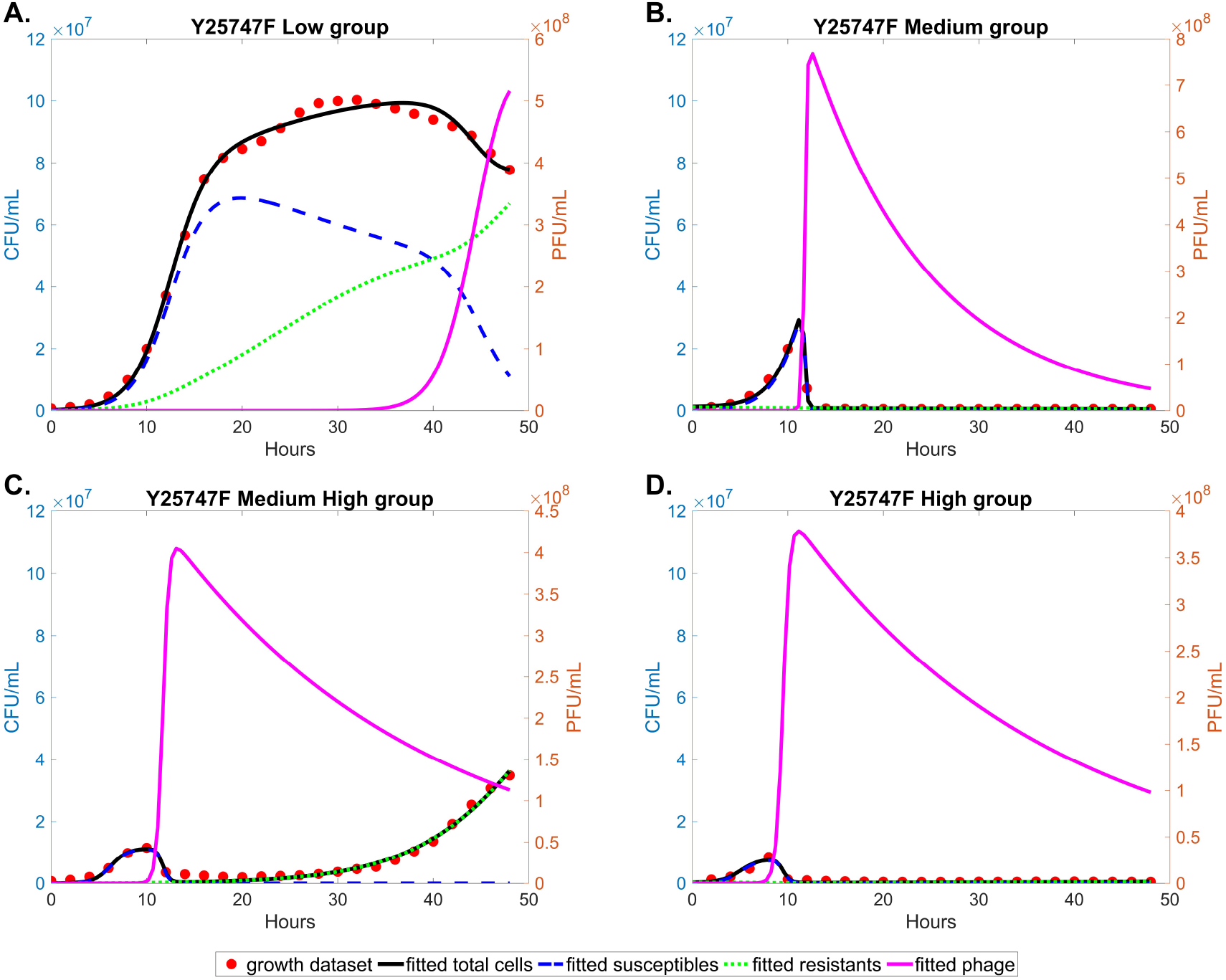
Dynamics of the mathematical model (1) when it is fitted to experimental growth dataset of strain 25747 Fern IDv1.

**Figure 7.**
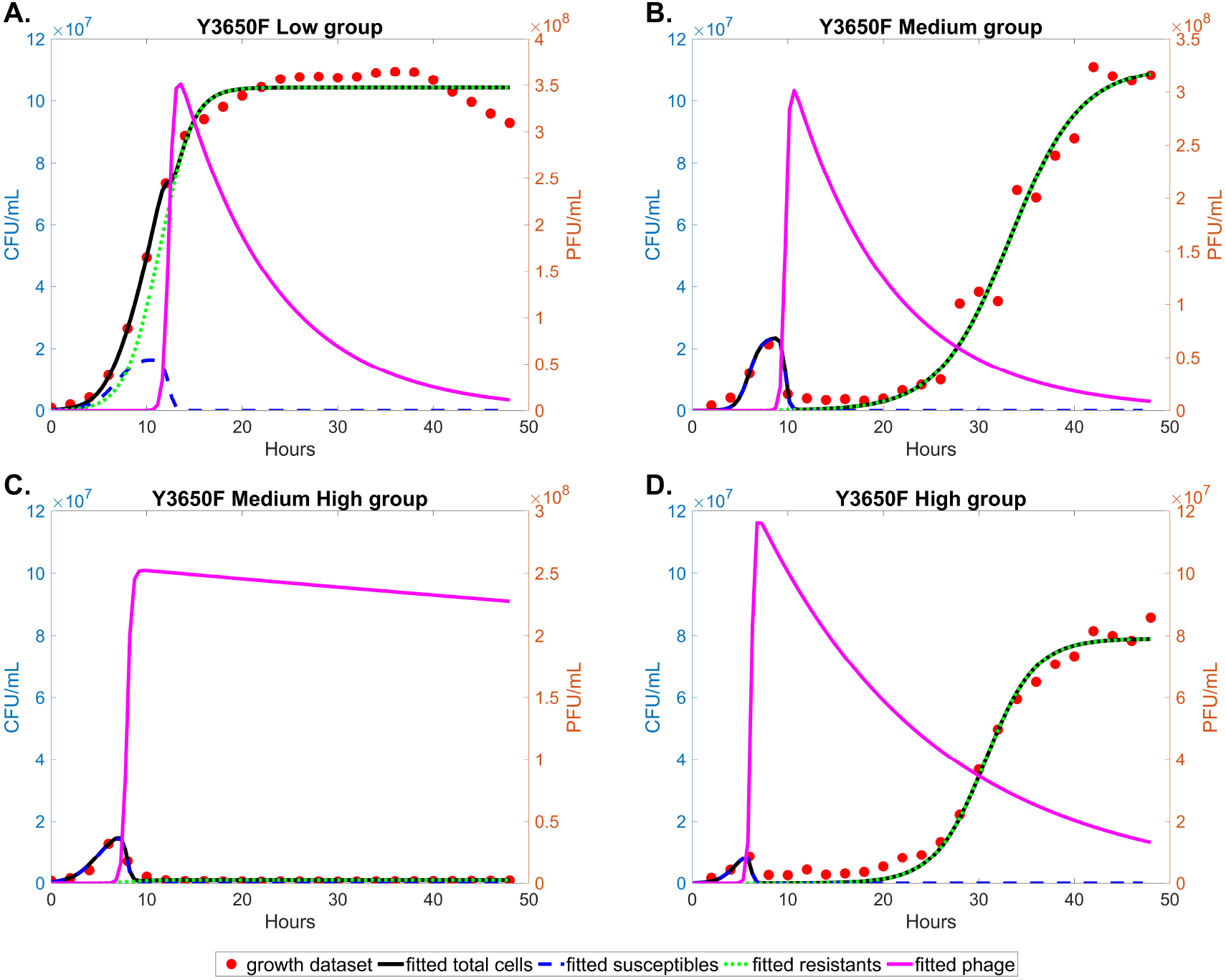
Dynamics of the mathematical model (1) when it is fitted to experimental growth dataset of strain Y-3650 Fern IDv1.

The dramatic, non-linear responses of competition parameters to increasing phage concentration revealed strain-specific competitive strategies that explain the observed growth patterns. PA103 and PAK both exhibited complex, multi-phase gap dynamics where |*a* − *a*_*rr*_| first narrowed at medium concentration before diverging at higher concentrations, suggesting initial competitive balance followed by asymmetric pressure. 25747 showed extreme gap widening at medium concentrations (driven by *a*_*rr*_) followed by narrowing at high concentrations, correlating with its unique ability to be completely suppressed even at medium phage levels. Y-3650 demonstrated the opposite pattern with gaps narrowing through medium-high concentrations before widening, reflecting its capacity for two-phase growth across a broader range of phage concentrations. These distinct parameter trajectories indicate that bacterial species employ fundamentally different competitive adaptations under phage stress.

Complete suppression patterns emerged through a critical interaction between proliferation thresholds and competitive dynamics that demonstrates the primacy of ecological context over intrinsic resistance mechanisms. Across all strains exhibiting complete suppression, only four core parameters (*r*, δ, *γ, ϕ*) maintained consistent resolution (Table 4), though additional parameters varied dramatically by strain. Examining Figure 3, the proliferation threshold *S*^∗^ showed complex, strain-specific responses to increasing phage concentration rather than simple monotonic decreases. For PA103, *S*^∗^ dropped sharply from Low to Medium concentrations, then stabilized through Medium High before decreasing further at High where suppression occurred. In contrast, PAK’s *S*^∗^ remained relatively stable across Low to Medium High before dropping at High concentration. 25747 demonstrated unique sensitivity, achieving complete suppression at Medium concentration despite *S*^∗^ values comparable to other strains. Y-3650’s *S*^∗^ fluctuated across concentrations, yet suppression occurred specifically at Medium High. These patterns reveal that suppression results not from *S*^∗^ alone but from its interaction with competitive asymmetries. When extreme gaps between competition parameters |*a* − *a*_*rr*_| coincided with specific *S*^∗^ ranges, bacterial populations could not recover. The low proliferation threshold creates conditions where phages maintain pressure even with minimal susceptible hosts, but this alone does not guarantee suppression—the competitive landscape must simultaneously prevent resistant bacteria from establishing viable populations. This pattern demonstrates that *S*^∗^ functions not merely as a kinetic parameter but as one component of a multi-dimensional ecological threshold that determines population fate [29, 30]

In contrast, two-phase growth patterns arose under diverse competitive conditions that varied dramatically across strains, challenging the notion of a single balanced dynamic. PA103 at Medium showed a moderate gap |*a* − *arr*| (~ 2.69) with susceptible bacteria exerting stronger interspecific competition than resistant bacteria experienced intraspecifically (*a > a*_*rr*_). PAK at Medium High exhibited an extreme gap (~ 42.2) following the same pattern. Intriguingly, 25747 at Medium High and Y-3650 at Medium showed the opposite arrangement with *a*_*rr*_ exceeding *a* (~ 8.82 and ~ 9.03 respectively), indicating that resistant bacteria faced stronger intraspecific competition. Most remarkably, Y-3650 demonstrated two-phase growth even at High phage concentration with an enormous gap (~ 456) where interspecific competition dominated. The observation that two-phase growth occurred across such varied competitive regimes—from *a*_*rr*_-dominated to ominated systems—suggests that the critical factor is not the balance of competition but rather the maintenance of conditions that allow temporal succession. Initially, susceptible bacteria grow rapidly when *a*_*ss*_ is moderate, then phages proliferate and suppress susceptibles, and finally resistant bacteria emerge when their competitive disadvantage (whether from high *a*_*rr*_ or high *a*) is offset by the removal of susceptible competitors by phages. These diverse pathways to two-phase dynamics suggest bacteria employ flexible resistance strategies—such as phase variation or biofilm formation—that provide temporary refugia without requiring specific competitive balance [31, 32]. The nearly identical resistance acquisition rates across these varied conditions confirms that two-phase growth emerges from ecological dynamics rather than mutational capacity [33].

The fitness cost parameter (*γ*) revealed unexpected complexity in Figure 3 that challenges conventional assumptions about resistance trade-offs. Rather than showing consistent costs across conditions, *γ* exhibited dramatic strain-specific and concentration-dependent variation. PA103 showed substantial fluctuation in fitness costs across phage concentrations, dropping from ~ 0.4 at Low to ~ 0.11 at Medium, rising to ~ 0.53 at Medium High, then decreasing to ~ 0.23 at High, suggesting dynamic modulation of resistance efficiency in response to varying selective pressures. PAK demonstrated a gradual decrease in fitness costs from ~ 0.7 at Low and Medium to ~ 0.5 at Medium High and ~ 0.33 at High, indicating progressive optimization of resistance mechanisms under increasing phage pressure. Most strikingly, 25747’s *γ* plummeted from 0.7 at Low to near zero (0.0009) at Medium—coinciding with complete suppression—before partially recovering at higher concentrations. Y-3650 showed more complex dynamics with *γ* varying between 0.17 and 0.99 across treatments. This heterogeneity indicates that fitness constraints are not fixed biological parameters but rather emerge from the interaction between resistance mechanisms and competitive environments. The observation that nearly identical resistance acquisition rates (δ remained relatively stable within each strain) produced vastly different fitness cost profiles suggests that bacteria can modulate the efficiency of resistance mechanisms depending on selective pressures, supporting recent findings on adaptive resistance strategies [34].

Our findings demonstrate that phage resistance evolution is fundamentally an ecological process where competitive interactions and fitness costs determine outcomes independently of intrinsic mutation rates. The consistent identifiability of four core parameters (*r*, δ, *γ, ϕ*) across all growth patterns and conditions establishes that resistance evolution follows predictable mathematical principles (Table 4, Figures 8-11). However, the dramatic variation in competitive parameters and their complex, non-linear responses to phage concentration (Figure 3) reveals that therapeutic success depends critically on controlling the competitive environment rather than suppressing mutation rates. The resistance acquisition rate δ remained relatively stable within strains across phage concentrations, yet produced completely different evolutionary outcomes—from successful resistance emergence to complete population collapse—depending solely on the competitive landscape [35]. These discoveries suggest that phage therapy strategies should focus on engineering competitive conditions that favor treatment success, with resistance management achieved through ecological manipulation rather than genetic intervention [36]. The strain-specific nature of competitive responses, particularly the distinct patterns between *P. aeruginosa* and *P. larvae* systems, indicates that therapeutic approaches must be tailored to the particular competitive strategies of target bacteria, emphasizing the importance of understanding bacterial ecology in clinical applications.

**Figure 8.**
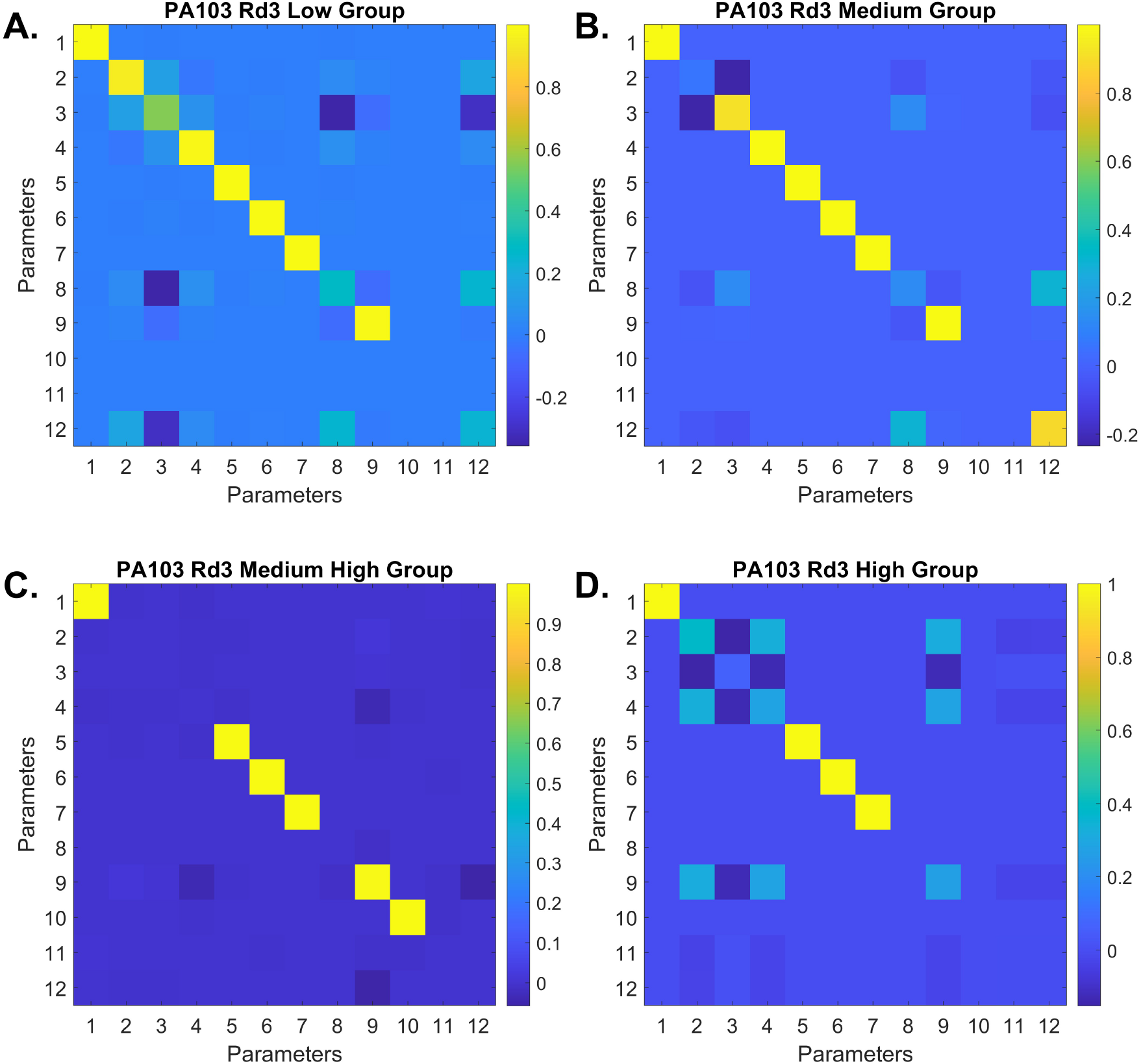
Model resolution matrices generated by fitted parameters in strain P. Aeruginosa PA103 treated by a cocktail of phages.

**Figure 9.**
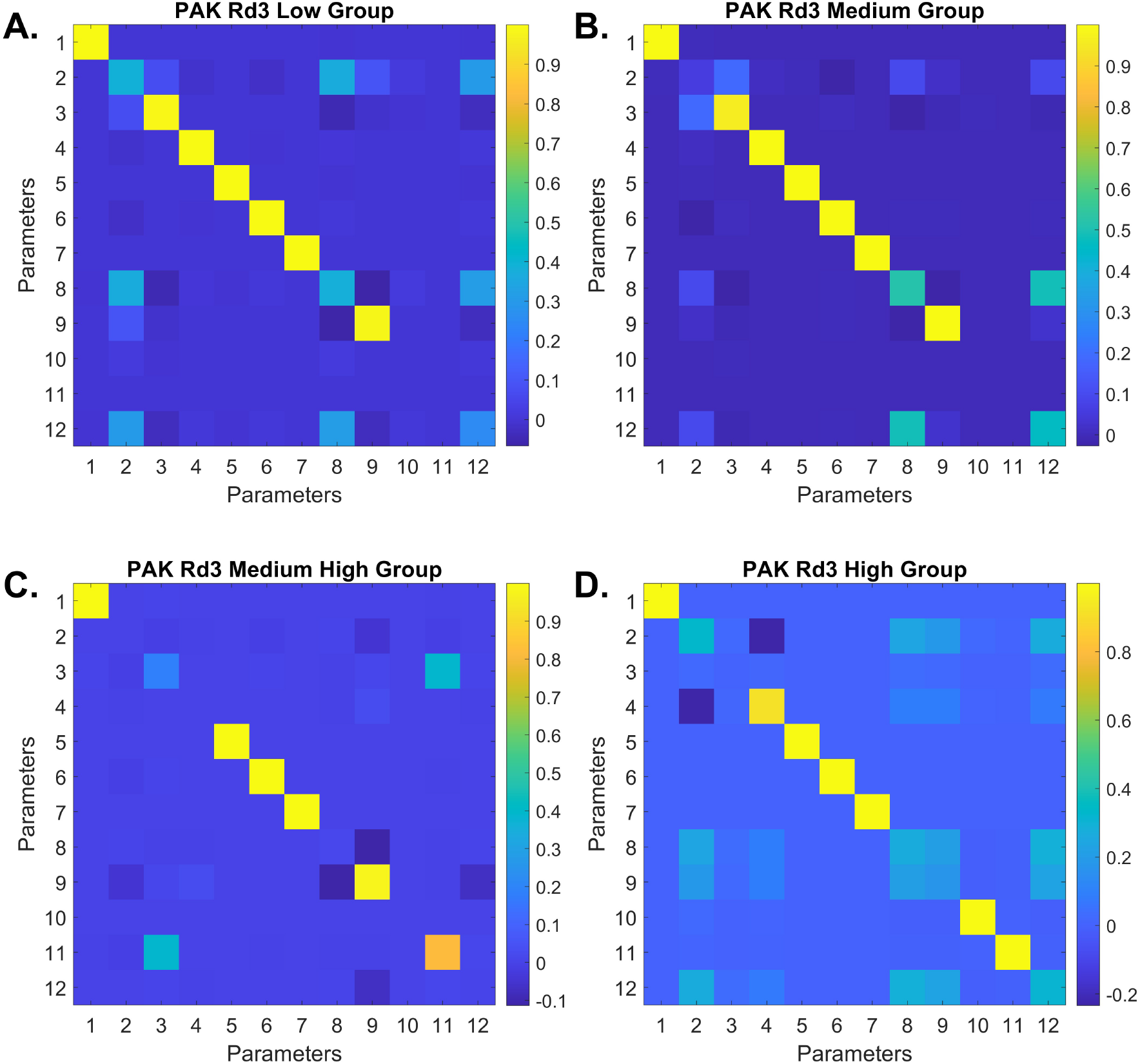
Model resolution matrices generated by fitted parameters in strain P. Aeruginosa PAK treated by a cocktail of phages.

**Figure 10.**
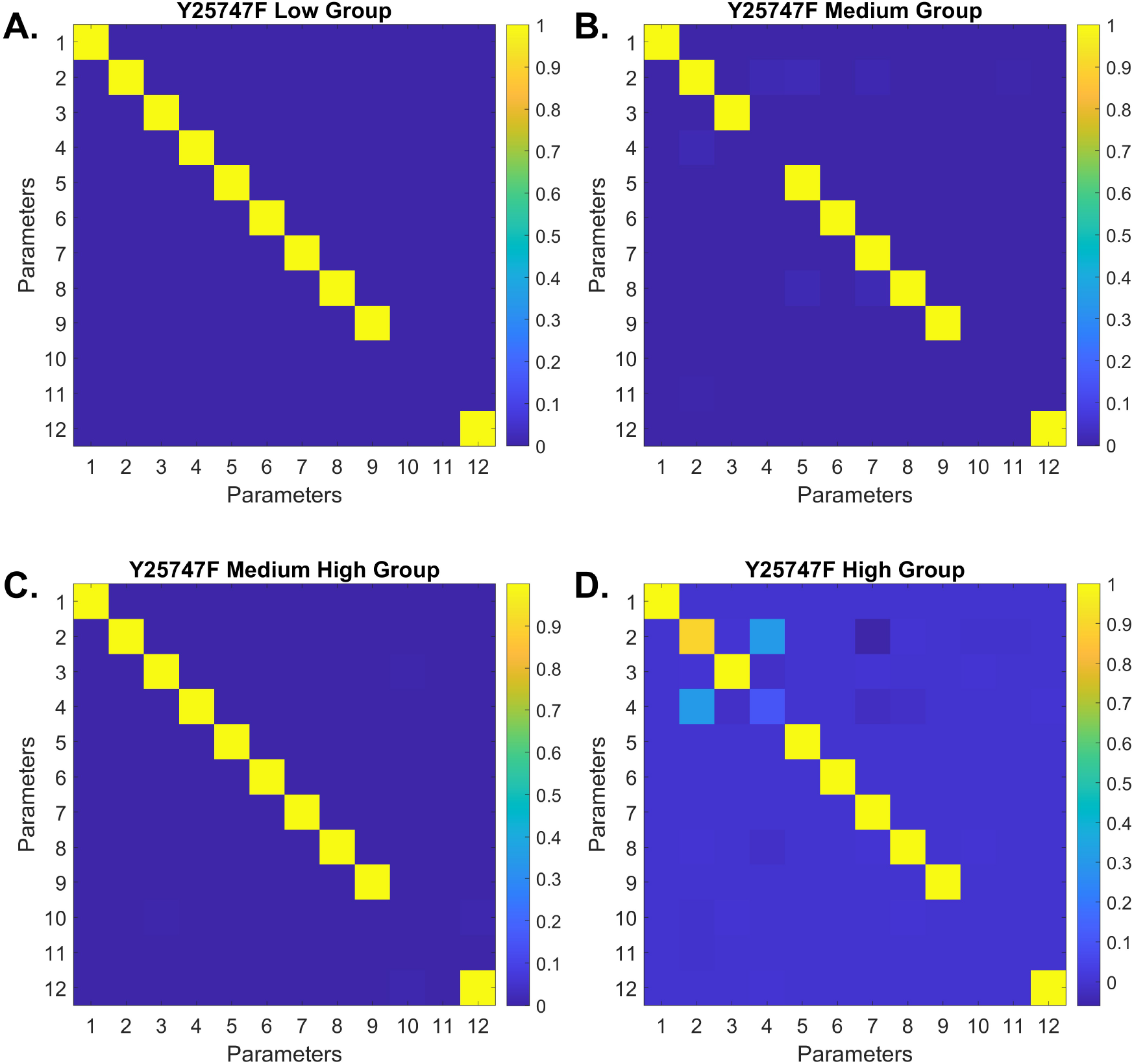
Model resolution matrices generated by fitted parameters in strain P. larvae 25747 treated by phage Fern IDv1.

**Figure 11.**
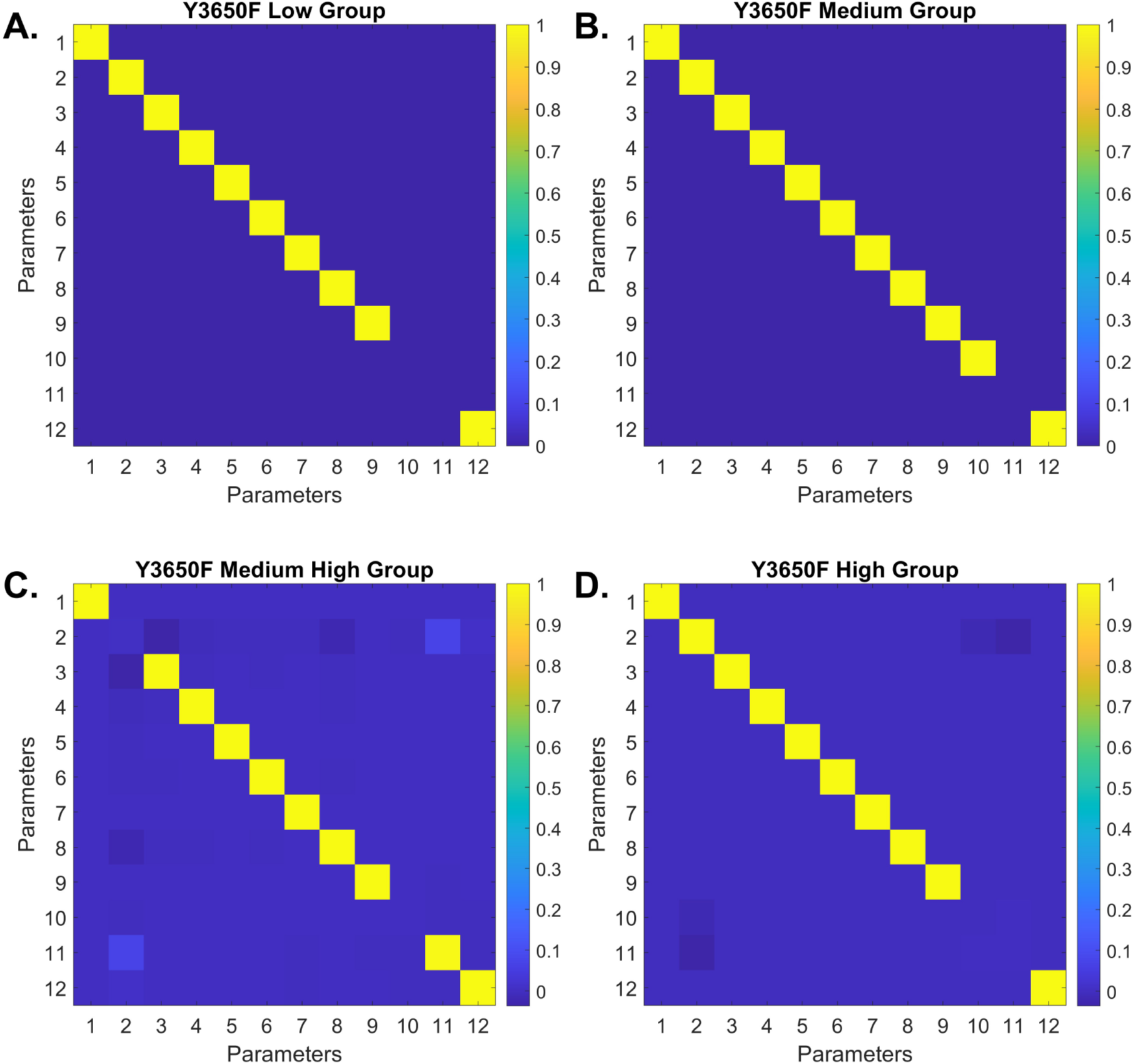
Model resolution matrices generated by fitted parameters in strain *P. larvae* Y-3650 treated by phage Fern IDv1.

## Acknowledgments

The research reported in this publication was supported by the National Institute of General Medical Sciences of the National Institutes of Health under Award Number P20GM104420 and the National Institute of Food and Agriculture of the US Department of Agriculture under Award Number 2023-67013-39067.

## Declarations

### Conflict of Interest

The authors declare no conflict of interest.

### Data availability

All experimental data analyzed and simulation MATLAB codes used to produce all the figures in this study are available at https://github.com/tphan86/BPI_model

## Authors’ contributions

- **Conceptualization**: JTVL
- **Methodology**: JTVL, AS, TP
- **Software**: TP
- **Data Collection**: AS, JS
- **Data analysis**: AS, JTVL
- **Mathematical Analysis and Fitting**: TP
- **Writing – original draft**: JTVL, TP
- **Writing – review & editing**: all authors
- **Visualization**: TP, JTVL
- **Project administration**: JTVL

## Notes

### Competing Interest Statement

The authors have declared no competing interest.

